# A putative ER-PM contact site complex relocalizes in response to Rare Earth Elements –induced endocytosis

**DOI:** 10.1101/720466

**Authors:** EunKyoung Lee, Brenda Vila Nova Santana, Elizabeth Samuels, Francisco Benitez-Fuente, Erica Corsi, Miguel A. Botella, Jessica Perez-Sancho, Steffen Vanneste, Jiří Friml, Alberto Macho, Aristea Alves Azevedo, Abel Rosado

## Abstract

In plant cells, environmental stressors induce changes in the cytosolic concentration of calcium ([Ca^2+^]_cyt_) that are transduced by Ca^2+^-sensing proteins. To confer specificity to the stress signaling response, [Ca^2+^]_cyt_ sensing must be tightly regulated in space and time; the molecular mechanisms that restrict the localization and dynamics of Ca^2+^ sensors in plants, however, are largely unknown. In this report, we identify a putative Ca^2+^-sensitive complex containing the synaptotagmins 1 and 5 (SYT1 and SYT5) and the Ca^2+^-dependent lipid binding protein (CLB1), which is enriched at ER-PM contact sites (EPCS) and relocalizes in response to Rare Earth Elements (REEs)-induced endocytosis. Our results show that endocytosed REEs influence cytosolic Ca^2+^ signaling, as indicated by the activation of the Ca^2+^/Calmodulin-based ratiometric sensor GCaMP3, and promote the cytoskeleton-dependent accumulation of ER-PM contact sites at the cell cortex. Based on these results, we propose that the EPCS-localized SYT1/SYT5/CLB1 complex is part of an evolutionarily conserved and spatially regulated Ca^2+^-responsive mechanism that control cER-PM communication during stress episodes.

## Introduction

Plant cells respond to different environmental stressors by changing their cytosolic free calcium concentration ([Ca^2+^]_cyt_). These changes in [Ca^2+^]_cyt_ act as secondary messengers, and their specific amplitude and duration (known as Ca^2+^ signature) provide specificity to the cellular responses **[1]**. In yeast and mammalian cells, the Endoplasmic Reticulum (ER) –Plasma Membrane (PM) Contact Sites’ (EPCS) microdomains act as Ca^2+^-sensitive platforms where EPCS-associated proteins sense changes in [Ca^2+^]_cyt_ and regulate the non-vesicular transfer of signaling molecules (e.g. lipids) between the cortical ER (cER) and the PM **[2-4]**. In Arabidopsis, the spatial and dynamic organization of EPCS is regulated by at least three families of EPCS components, namely, synaptotagmins (SYTs), Vesicle-associated membrane protein (VAMP)-associated proteins 27 (VAP27s), and VAP-RELATED SUPPRESSORS OF TOO MANY MOUTHS (VSTs) **[5]**. These EPCS components in plants serve a number of well-characterized functions including response to biotic and abiotic stressors, **[6-14]**, the control of the interactions between the ER and the cortical cytoskeleton **[15-16]**, and the activation of signal transduction events through the activation of Receptor–like Kinases at the PM **[17]**. Despite this wealth of functional information, the specific responses of plant EPCS components to stress-derived Ca^2+^ signals remain largely uncharacterized.

This work focuses on the Ca^2+^-dependent phospholipid binding protein SYT1, which shares structural organization with the mammalian extended-synaptotagmins (E-Syts) and yeast tricalbins (Tcbs) families of Ca^2+^-responsive EPCS tethers **[18]**. At the structural level, SYT1 is anchored to the cER, targeted to EPCS through a putative synaptotagmin-like mitochondrial lipid-binding domain (SMP) **[19]**, and docked to the PM through Ca^2+^-dependent interactions between its C2 domains and negatively charged phospholipids **[6, 7, 20, 21]**. At the functional level, SYT1 interacts with elements of the exocytotic soluble N-ethylmaleimide-sensitive factor attachment protein receptor complexes (SNAREs) **[9]**, phytosterol binding proteins **[24]**, and reticulon proteins **[25]**. These interactions are required for the maintenance of the cER stability **[13]**, the control of immune secretory pathways **[9]**, the regulation of cell-to-cell communication **[10-12]**, and the tolerance to ionic, mechanical and freezing stresses **[6, 7, 14]**. The specific role of stress-derived Ca^2+^ signals in the regulation of SYT1 activity and EPCS organization, however, remains largely uncharacterized.

This study identifies a putative tethering complex formed by the Synaptotagmins 1 (SYT1) and 5 (SYT5) and the Ca^2+^-dependent lipid binding protein CLB1. This complex in enriched at ER-PM contact sites, responds to environmental stresses, and induces EPCS rearrangements in response to Rare Earth Elements (REEs)-induced endocytosis. Mechanistically, this study shows that long-term exposure to REEs induces the accumulation of phosphatidylinositol-4-phosphate (PI4P) at the PM, promotes the internalization of PI4P-containing vesicles, activates the Calmodulin-GFP based Ca^2+^ sensor GCaMP3 **[26]**, and triggers the concomitant accumulation of SYT1/SYT5/CLB1 containing EPCS at the cell cortex. The study also highlights clear differences between the EPCS remodeling triggered by NaCl, which is largely cytoskeleton-independent **[8]**, and the EPCS accumulation triggered by REE-endocytosis, which is largely cytoskeleton-dependent. Our results support a model where EPCS tethering complexes act as Ca^2+^ sensing platforms that regulate cortical ER-PM communication in response to stress-induced changes in cytosolic Ca^2+^ signaling. These examples illustrate the plasticity that governs cER-PM communication, an evolutive milestone of eukaryotic organisms, and highlight the role of evolutionarily conserved tethering complexes in the coordinatation of cellular responses to environmental stress.

## Results

### Identification of SYT1/SYT5/CLB1 as a putative Ca^2+^-binding tethering complex in Arabidopsis

SYT1 is a protein tether implicated in the establishment, organization, and function of plant ER-PM contact sites (EPCS) **[6-14]**. Because the SYT1 orthologs in mammals and yeast establish tethering complexes *in vivo* **[27-28]**, and multiple proteins containing C2 domains and TM regions are able to tether membranes in Arabidopsis plasmodesmata **[29]**, we searched for additional proteins physically associated with SYT1 that could participate in EPCS establishment and/or regulation. For this purpose, we used a SYT1-GFP line in the *syt1-2* background **[8]** and performed immunoprecipitation (IP) assays using agarose beads coupled to an anti-GFP nanobody (GFP-Trap beads). The IP results from three independent biological replicates provided a large number of proteins physically associated with SYT1 that we identified using liquid chromatography coupled to tandem mass-spectrometry (LC-MS/MS). We filtered the results using the following criteria: 1. Presence in all three biological replicates; 2. Detection of two or more exclusive unique peptides; and 3. Absence in the negative immunoprecipitation control (IP using a transgenic line expressing free GFP). After applying these filters we identified two putative SYT1 interactors: the Arabidopsis Synaptotagmin 5 (*At1g05500*, SYT5), an uncharacterized member of the Arabidopsis synaptotagmin family, and the Ca^2+^ and lipid binding protein 1 (*At3g61050*, CLB1), a protein involved in drought and NaCl stress responses **[30] (Table 1A and Table S1)**. To confirm the interaction between SYT1 and these two proteins we used a targeted co-IP assay. For this experiment we generated a transgenic line expressing SYT5-GFP under its native promoter (SYT5::SYT5-GFP) and a transgenic line expressing CLB1-GFP under a constitutive ubiquitin 10 promoter (*pUB10::CLB1-GFP*). Upon immunoprecipitation using SYT5::SYT5-GFP and *pUB10::CLB1-GFP*, we performed hybridization with a custom antibody raised against SYT1 [7]. Our results confirmed that the native SYT1 associates with SYT5-GFP and CLB1-GFP, but not with free GFP **(Figure 1B)**.

**Figure 1.**
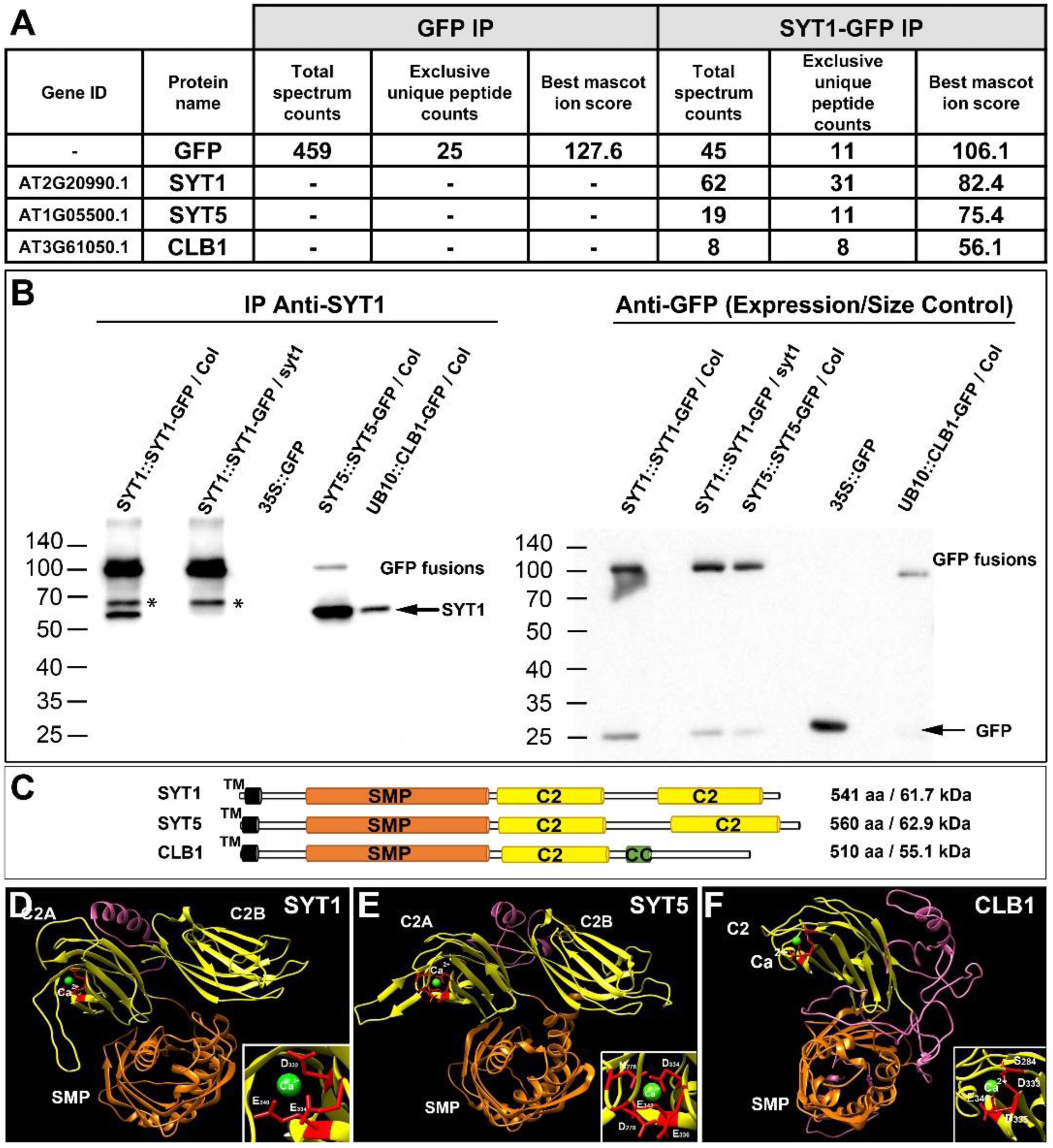
The Ca^2+^-dependent phospholipid binding proteins SYT5 and CLB1 interact with SYT1. Table 1A. Peptide counts detected upon GFP immunoprecipitation followed by LC-MS/MS analysis using Arabidopsis plants expressing GFP (control) and SYT1-GFP. Numbers indicate the total spectrum counts corresponding to the indicated proteins, and the exclusive unique peptides represented within them. The best mascot ion score among these peptides is indicated. The number of peptides corresponding to GFP is shown for reference. This result is representative from 3 independent experiments (For details on the replicates see **Table S1**). **B)** Arabidopsis transgenic plants expressing SYT1-GFP, SYT5-GFP and CLB1-GFP were used for immunoprecipitation using GFP-trap beads. Plants expressing free GFP were used as control. The immunoprecipitated proteins were separated by SDS-PAGE, and western blots were analysed using anti-SYT1 (left panel) or anti-GFP (right panel) antibodies. Molecular weight (kDa) marker bands are indicated for reference. The arrow indicates the expected MW for SYT1.*indicate a SYT1-GFP derived fragment recognized by the SYT1 antibody. **C)** Schematic representation of the functional domains of SYT1, SYT5 and CLB1. TM: Transmembrane Domain, SMP: synaptotagmin-like mitochondrial-lipid binding domain, C2: phospholipid binding domains, CC: Coiled coil domain. **D)** 3D structures and Ca^2+^ binding sites in the predicted cytosolic regions of SYT1, SYT5 and CLB1 identified using Phyre2 and 3DLigand site. Important aminoacid residues for Ca^2+^ binding are indicated in red.

To determine the relative abundance and tissue specificity of the putative <u>S</u>YT1/<u>S</u>YT5/<u>C</u>LB1 complex (hereafter SSC complex) we used freely available RNAseq data from the transcriptomic variation database (TraVa) **[31]. Table S2** shows that the SYT1, SYT5 and CLB1 genes are ubiquitously expressed, and that the *SYT1* transcripts are, on average, three times more abundant than *SYT5’*s and two times more abundant than *CLB1*’s across multiple tissues. We also assigned putative functions for SYT5 and CLB1 through the identification of their functional domains using Pfam **[32]** and TMHMM2 **[33]** databases. The analyses show that SYT1, SYT5, and CLB1 share a common domain architecture comprising a putative single N-terminal TM domain, a ∼40 amino acid linker, a cytoplasm-exposed synaptotagmin-like mitochondrial lipid-binding protein (SMP) domain; and one (CLB1) or two (SYT1 and SYT5) phospholipid-binding C2 domains harboring Lysine/Arginine rich (K/R-rich) polybasic patches **(Figure 1C and Figure S1)**. Next, we generated 3D structures for the predicted cytosolic regions of these proteins (SYT1_30-541_), (SYT5_23-560_) and (CLB1_22-510_) using the Phyre2 **[34]** and 3DLigandSite **[35]** modelling tools. The results show that 89% of the SYT1 aminoacid residues, 87% of the SYT5 aminoacid residues, and 69% of the CLB1 residues could be modelled with >90% confidence using the crystal structure of the mammalian extended synaptotagmin 2 as a template **(Figures 1D-1F)**. Furthermore, the 3DLigandSite analyses also show that all three proteins contain a single Ca^2+^ binding site, whose position was determined with confidence levels above the 99% threshold **(Figures 1D-1F Insets)**. Based on the *in vitro* interactions, and the subsequent bioinformatics analyses, we propose that SYT1, SYT5 and CLB1 are part of a putative Ca^2+^-binding tethering complex (hereafter SSC complex) that is ubiquitously expressed in Arabidopsis.

### The putative SSC complex is enriched at EPCS and is required for root hair polarity maintenance

To establish unequivocally the subcellular localization of the putative SSC complex *in vivo*, we generated fluorescent SYT5-GFP and CLB1-GFP marker lines driven by their respective endogenous promoters, and used confocal microscopy to compare the SYT5-GFP and CLB1-GFP subcellular localization with that of the SYT1-GFP marker **[8]. Figure 2** shows that the SYT5-GFP and CLB1-GFP localization strongly resemble that of the SYT1-GFP marker in all tissues analyzed. These localizations include a “beads and strings” arrangement in cotyledon epidermal cells **(Figures 2A-2C)**, perinuclear labelling consistent with ER in root meristematic cells **(Figures 2D-2F)**, strong accumulation of signals at root hair initiation sites **(Figures 2G-2I)**, and associations to the cell wall through Hechtian strands **(Figures 2J-2L)**.

**Figure 2.**
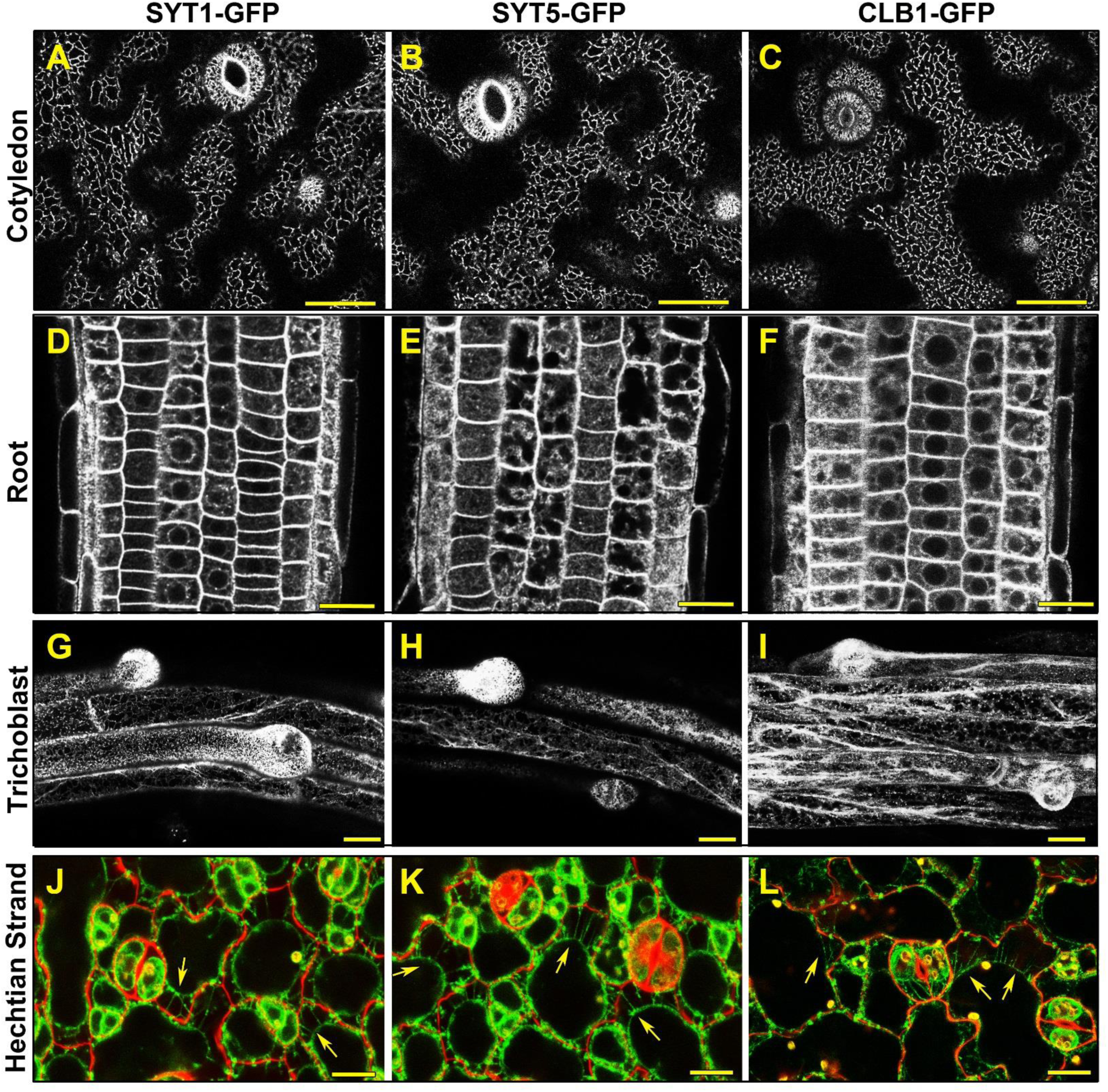
Subcellular localization of the putative SYT1/SYT5/CLB1 tethering complex. **(A-C)** Subcellular localization of the SYT1-GFP and SYT5-GFP and CLB1-GFP markers in epidermal cells of 5-d-old cotyledons **(A-C)**, root meristematic cells of 5-d-old seedlings **(D-F)**, and emerging root hairs in 7-d-old seedlings **(G-I)** Scale bars (A-I) = 25 μm. **(J-L)** SYT1-GFP, SYT5-GFP and CLB1-GFP are present in Hechtian strands. 5-d-old cotyledon epidermal cells expressing the different markers were plasmolyzed for 4 h using 0.4 M mannitol. The images are an overlay of propidium iodide stained cell walls with the localization of the GFP fusion proteins in green Arrows indicate Hechtian strands. Scale bar = 20 μm.

Given that SYT1, SYT5, and CLB1 have similar functional domains and subcellular localization, we asked whether the NaCl hypersensitivity previously described for the *syt1-2* mutant **[6]** was a common feature shared by the *syt5* and/or the *clb1* loss-of function mutants. To answer this question, we isolated two homozygous lines harboring T-DNA insertions in the seventh intron of the *At1g05500* locus (*syt5-1*, SALK_036961) and the eighth intron of the *At3g61050* locus (*clb1-2*, SALK_006298) **(Figure S2A)**. Once we confirmed that *SYT5* and *CLB1* transcripts were not present by qRTPCR **(Figure S2B)**, we generated the triple *syt1-2/ syt5-1 /clb1-2* mutant and performed root elongation assays in the presence of NaCl as described in **[6]. Figure 3A** shows that, in the presence of NaCl, *syt5-1 and clb1-2* root elongation is indistinguishable from that of the wild type (WT) and that *syt1-2* displays a hypersensitive response similar to that of the triple *syt1-2/ syt5-1 /clb1-2* mutant. Remarkably, the NaCl treatments also induced root hair polar growth defects in *syt1-2* and *syt5-1* but not *clb1-2* backgrounds. These defects characterized by root hair skewing and branching, were more severe in the triple *syt1-2/ syt5-1 /clb1-2* mutant **(Figure 3B)**. Combined, these results suggest that SYT1 is a housekeeping gene required for proper root elongation during NaCl stress, and that the activity of additional components of the SSC complex is required for proper root hair polarization maintenance.

**Figure 3.**
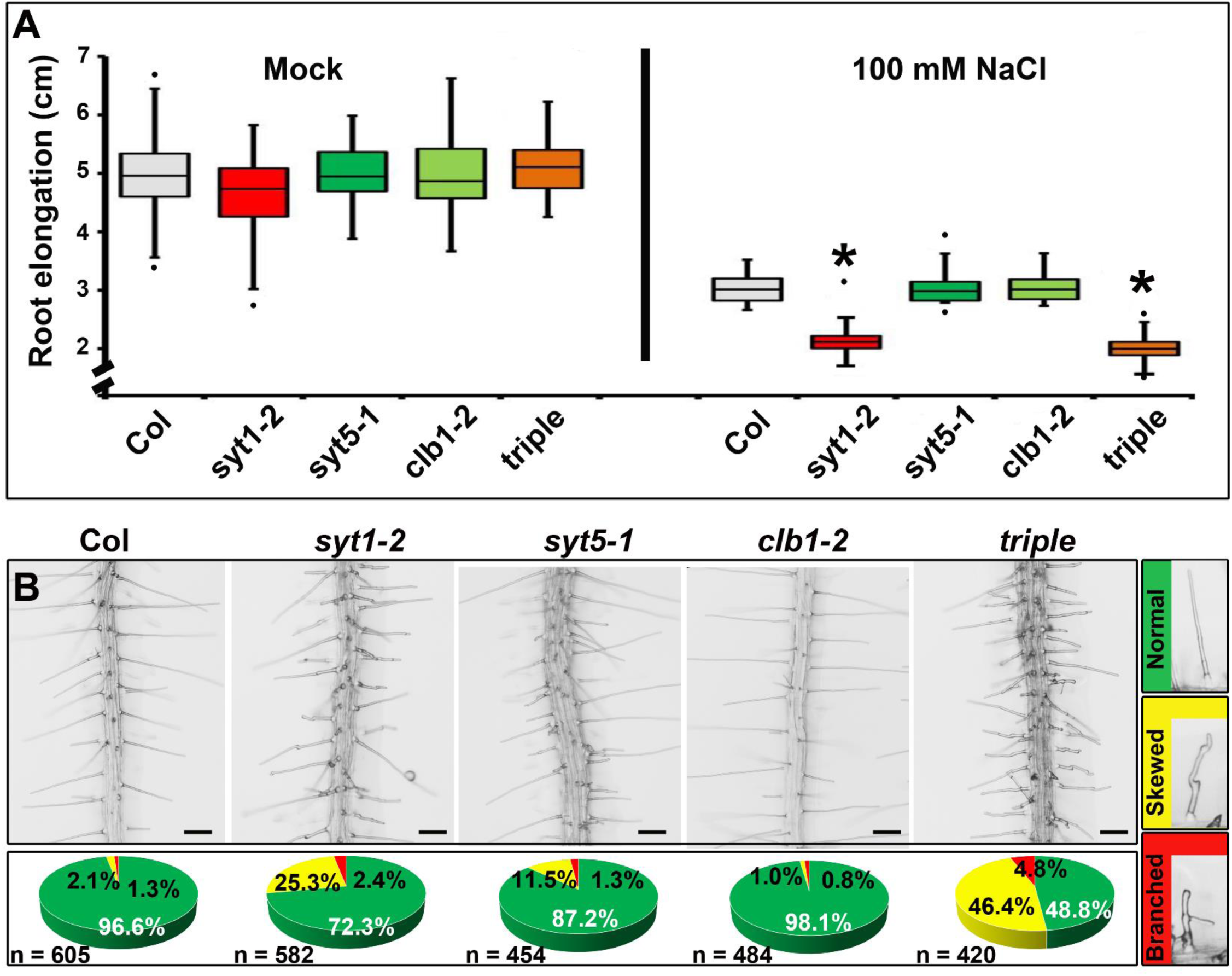
Responses to NaCl stress. **(A)** Root elongation NaCl dosage analysis. Seedlings were grown in one tenth-strength MS medium for 4 d and then transferred to the same media supplemented with the NaCl concentrations indicated. n = 60 seedlings / treatment. Asterisks indicate significant differences between the wild type (Col) and the mutants (P < 0.05) **(B)** Representative roots from WT (Col), *syt1-2, syt5-1, clb1-2* single and *syt1-2 syt5-1 clb1-2* triple mutants showing the root hair morphology upon NaCl stress. *Arabidopsis* seedlings were grown vertically for 5 d on one tenth-strength MS medium supplemented with 50 mM NaCl and directly imaged on the growth media without mounting. **(C)** Quantification of root hair phenotypes. Right panels show representative root hair morphologies categorized as normal (green), skewed (yellow) or branched (red). Data represent the percentage of root hairs on each category and n represents the number of root hairs quantified for each genotype. Scale bars = 200 μm.

### The putative SSC complex relocalizes in response to treatments with Rare Earth Elements

The known effects of the *syt1-2* mutation in Ca^2+^-regulated stress responses **[6, 7, 14]**, the observed root hair polar growth defects that can be associated to aberrant [Ca^2+^]_cyt_ oscillations **[36]**, and the identification of a highly conserved Ca^2+^ binding site in SYT1, SYT5 and CLB1 **(Figure 1D-1F insets)** prompted us to ask whether Ca^2+^ signals could influence the localization and dynamics of the SSC complex and induce changes in EPCS organization. To address this question, we first analyzed the effect of extracellular Ca^2+^ depletion on SYT1-GFP, SYT5-GFP and CLB1-GFP localization using the extracellular Ca^2+^ chelating agents ethylene glycol-bis(β-aminoethylether) – β N,N,N’,N’-tetraacetic acid (EGTA) and bis-(*o*-aminophenoxy) ethane-*N,N,N’,N’*-tetra-acetic acid (BAPTA) **[37, 38]**. Our results show that the long-term depletion of free apoplastic Ca^2+^ induced by either EGTA or BAPTA did not have a significant effect in the number of SYT1-GFP, SYT5-GFP labelled “beads” at the cell cortex, and did not induce gross morphological changes in the cER as indicated by the GFP-HDEL marker **(Figures 4A-4I and Figures 4P-4R)**. In sharp contrast, the blockage of Ca^2+^ entry using the Rare Earth Elements (REEs) La^3+^ and Gd^3+^ as non-selective cation channel blockers **[39-42]** induced severe changes in both the localization and abundance of componentsof the SSC complex **(Figures 4J – 4O)**. Treatments with REEs caused a 4 to 5 fold increase in the number of SYT1-GFP and SYT5-GFP labelled “beads” at the cell cortex **(Figures 4P-4Q)**, which likely represent newly established EPCS, as indicated by the increase in the number of beads labelled by the artificial EPCS marker MAPPER-GFP **[8] (Figure S3)**. This increase in EPCS number was accompanied by cER rearrangements characterized by a significant reduction of the average reticule size of the cER network **(Figure 4R)**. Remarkably, treatments with EGTA and BAPTA did not change the abundance and distribution of the CLB1-GFP marker **(Figures S4A-S4C)**, but treatments with REEs often caused a significant reduction in the CLB1-GFP fluorescent signal that hindered our ability to further analyze the subsequent CLB1 responses **(Figures S4D-S4E)**.

**Figure 4.**
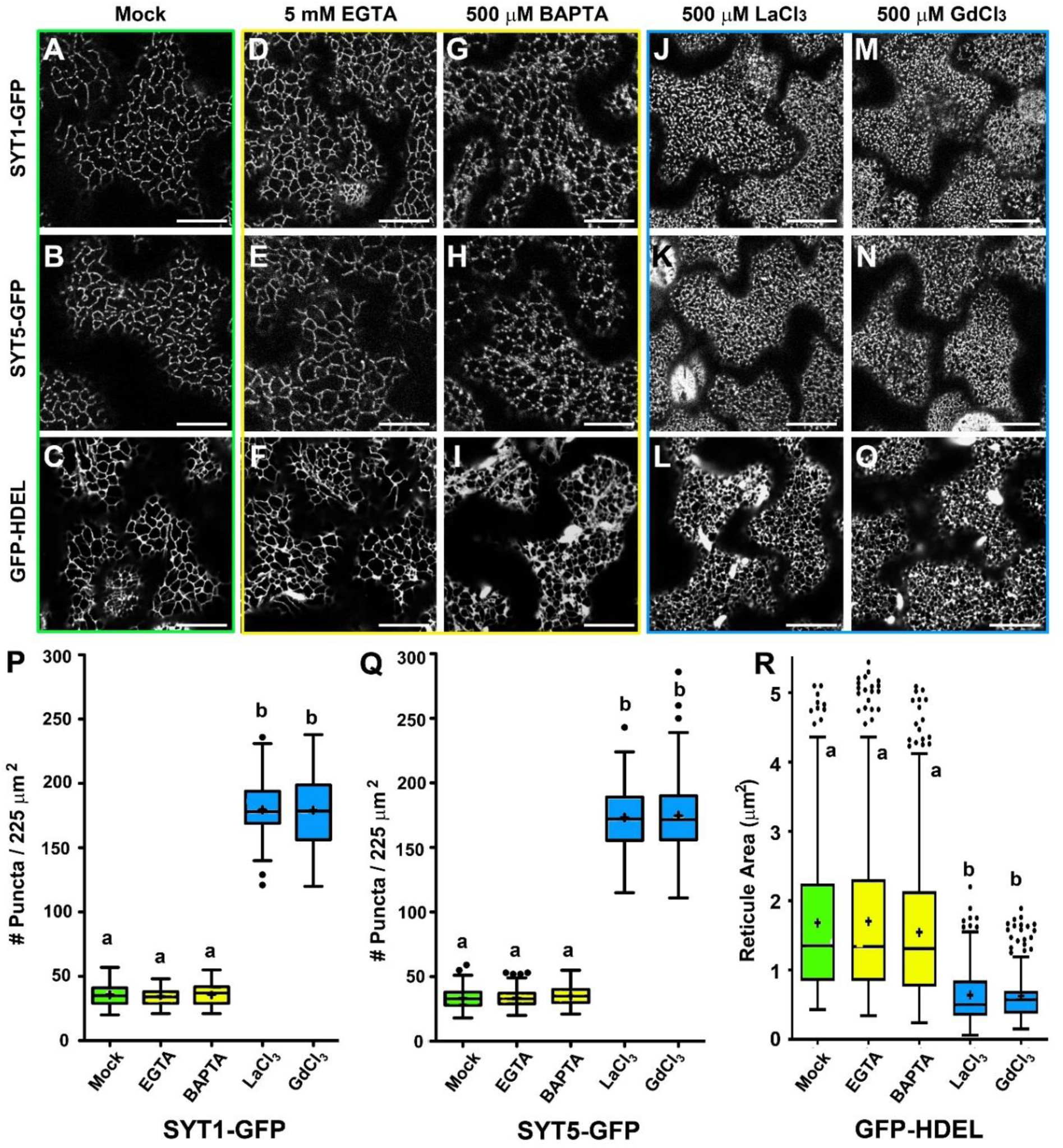
Treatments with non-selective Ca^2+^ channel blockers increase the number of putative SSC complexes at the cell cortex. 5-d-old SYT1-GFP, SYT5-GFP, and HDEL-GFP seedlings were treated in liquid 1/10 strength MS medium supplemented with Mock **(A-C)** EGTA (5 mM, 2 h) **(D-F)**, BAPTA (250 µM, 2 h) **(G-I)**, LaCl3 (500 µM / 16 h) **(J-L)** or GdCl3 (500 µM / 16 h) **(M-O**) prior to imaging and quantification. **(P-Q)** Quantification of SYT1-GFP and SYT5-GFP puncta at the cell cortex upon chemical treatments. The number of SYT1-GFP and SYT5-GFP puncta were scored from 50-60 arbitrary 225 mm^2^ ROIs from at least 15 cells from 5 different seedlings. **R)** Quantification of HDEL-GFP reticulation. For each treatment the number of closed reticules was scored using 50 – 60 arbitrary 225 mm^2^ ROIs from at least 15 cells from 5 independent seedlings. In the box and whiskers plots, the center line represents the median number of closed reticules / 225 mm^2^, the top and bottom edges are the 25th and 75th percentiles of the distribution, and the ends of the whiskers are set at 1.5 times the IQR. When present, the minimum / maximum values outside the IQR are shown as outliers (dots). Letters indicate statistically significant differences using Tukey multiple pairwise-comparisons p <0.05 Scale bars = 20 μm

### REE-endocytosis activates Ca^2+^ signaling and induces cytoskeleton-dependent changes in SYT1-GFP and SYT5-GFP localization

Previous reports have shown that long-term treatments with REEs induce endocytosis and promote the release of REEs to the cytosol **[43, 44]**, so we asked whether REE-endocytosis underlies the observed changes in SYT1-GFP and SYT5-GFP localization following such a treatment. **Figures 5A-5C** show that the REE treatments promote the accumulation and internalization of phosphatidylinositol-4-phosphate (PI4P) containing-vesicles, as indicated by the ratiometric sensor citrine 1xPH^FAPP1^ **[45]**. These vesicles are likely to be endocytic as indicated by their co-localization with the FM4-64 dye in short-term (3 minutes) treatments **(Figure S5)**. In agreement with chemical studies showing that REEs form stable complexes with polydentate chelators **[46]**, the addition of EGTA to growth media was sufficient to inhibit the REE-induced endocytosis **(Figures 5D-5F and Figure 5G)**, reverse the SYT1-GFP localization changes associated to REE internalization **(Figure S6)**, and reduce the overall REE toxicity in long-term treated seedlings **(Figure S7)**. To understand the mechanisms underlying the change in SYT1-GFP and SYT5-GFP localization in response to REE-induced endocytosis, we assesed whether endocytosed REEs could act as Ca^2+^ surrogates by activating Ca^2+^ signaling in the cytosol. To answer this question, we analyzed the effect of endocytosed REEs in the activity of the calmodulin-based ratiometric Ca^2+^ sensor GCaMP3 **[26]. Figures 5H-5M** and **Figure 5N** show that long-term REE treatments (16 h) induced a 2-3 fold increase in the GCaMP3 sensor signal that was abolished by the addition of the extracellular chelator EGTA. This result confirms previous biochemical and physiological studies showing that REEs can interact with Calmodulins **[47-49]**.

**Figure 5.**
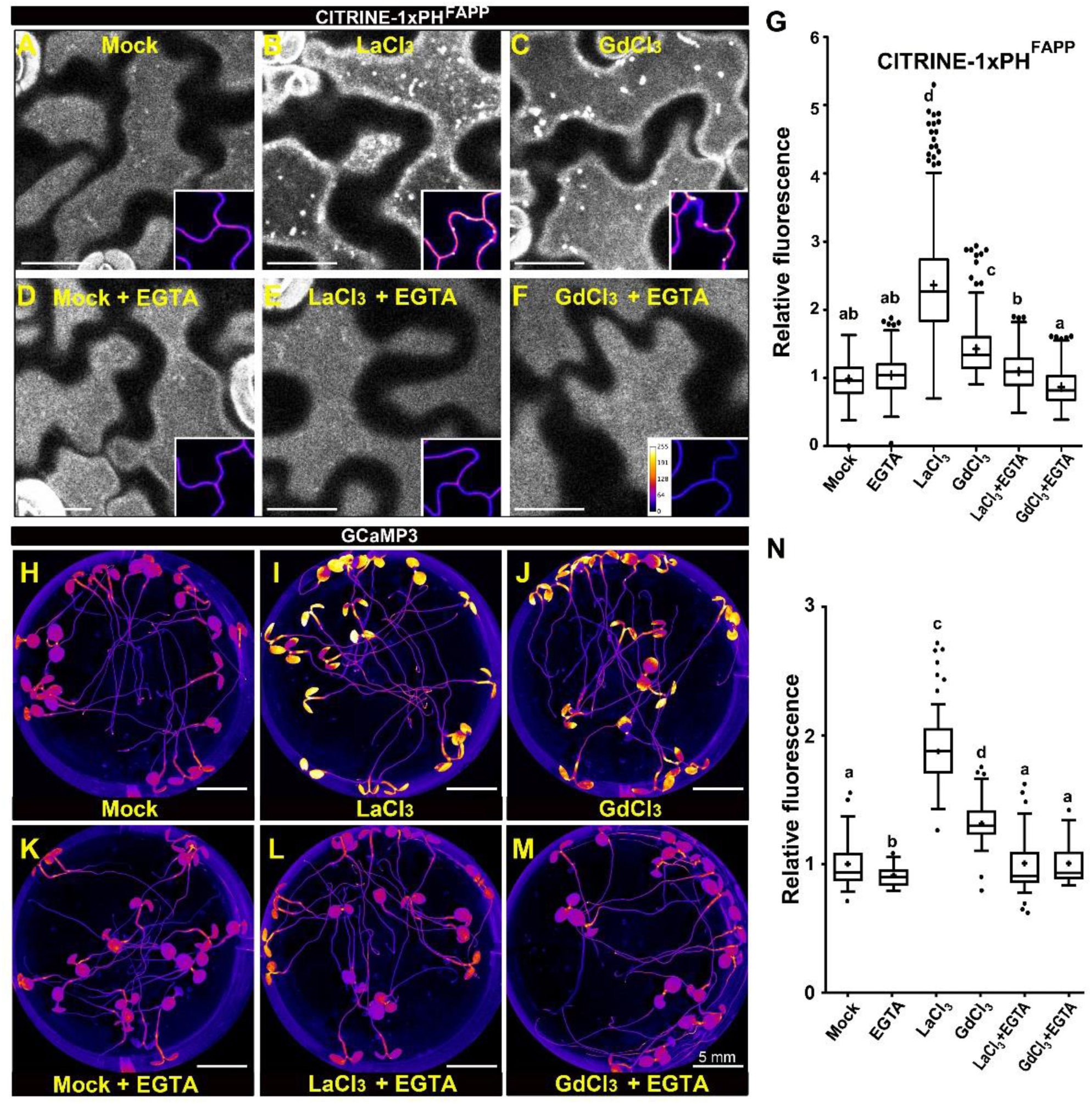
REE induces the accumulation and internalization of PI4P-containing vesicles and activates the GCaMP3 sensor. **(A-F)** Confocal images of the cell cortex in cotyledon epidermal cells expressing the CITRINE-1xPH^FAPP^ marker. 5-d-old seedlings were treated in liquid one-tenth-strength MS medium supplemented with Mock, 16 h **(A)**, LaCl3 (500 µM / 16 h) **(B)**, GdCl3 (500 µM / 16 h) **(C)**, or the same media supplemented with 5 mM EGTA **(D-F)**. The presence of endocytic vesicles is shown as bright dots at the cell cortex, and PI4P accumulation is shown in the insets as color-coded pixel intensity following the LUT scale shown in F. **G)** Quantification of the CITRINE-tagged 1xPH^FAPP^ signal relative to mock conditions. The letters indicate statistically significant differences using Tukey multiple pairwise-comparisons p <0.05. Scale bars = 20 mm. (H-M) Fluorescence images of seedlings expressing the GCaMP3 Ca^2+^ sensor. 5-d-old seedlings were treated in liquid one-tenth-strength MS medium supplemented with Mock, 16 h **(H)**, LaCl3 (500 µM / 16 h) **(I)**, GdCl3 (500 µM / 16 h) **(J)**, or the same media supplemented with 5 mM EGTA **(K-M)**. The activity of the Ca^2+^ sensor is shown as color-coded pixel intensity following the LUT scale shown in F. **N)** Quantification of the GCaMP3 signal relative to mock conditions. Scale bars = 5 mm In the G and N plots, the center line represents the median fluorescence intensity fold increase relative to mock, the cross represents the mean fluorescent intensity, the top and bottom edges are the 25^th^ and 75^th^ percentiles of the distribution, and the ends of the whiskers are set at 1.5 times the interquartile range (IQR). All values outside the IQR are shown as outliers. At least 100 regions of interest (ROIs) **(G)** or 50 seedlings **(N)** were measured for each treatment.

Because the delivery of SYT1-GFP to EPCS is cytoskeleton dependent but its rearrangement in response to NaCl is cytoskeleton independent **[8]**, we asked whether the actin cytoskeleton and/or the microtubule network play a role in the REE-induced SYT1-GFP and SYT5-GFP relocalization process. **Figure S8** shows that, different thanthe previously described NaCl treatment**[8]**, the treatment with LaCl_3_ induces SYT1-GFP and SYT5-GFP relocalization without visible disruption of thethe cortical cytoskeleton organization. Furthermore, our results show that the effect of LaCl_3_ in SYT1-GFP and SYT5-GFP localization is partially abolished by pre-treatments with the cytoskeleton depolymerizing drugs Oryzalin and Latrunculin B **(Figure 6**). Together, these results suggest that a cytoskeleton-dependent remodeling mechanism underlies the changes in SYT1-GFP and SYT5-GFP localization in response to LaCl_3_ treatments **(Figure 7)**.

**Figure 6.**
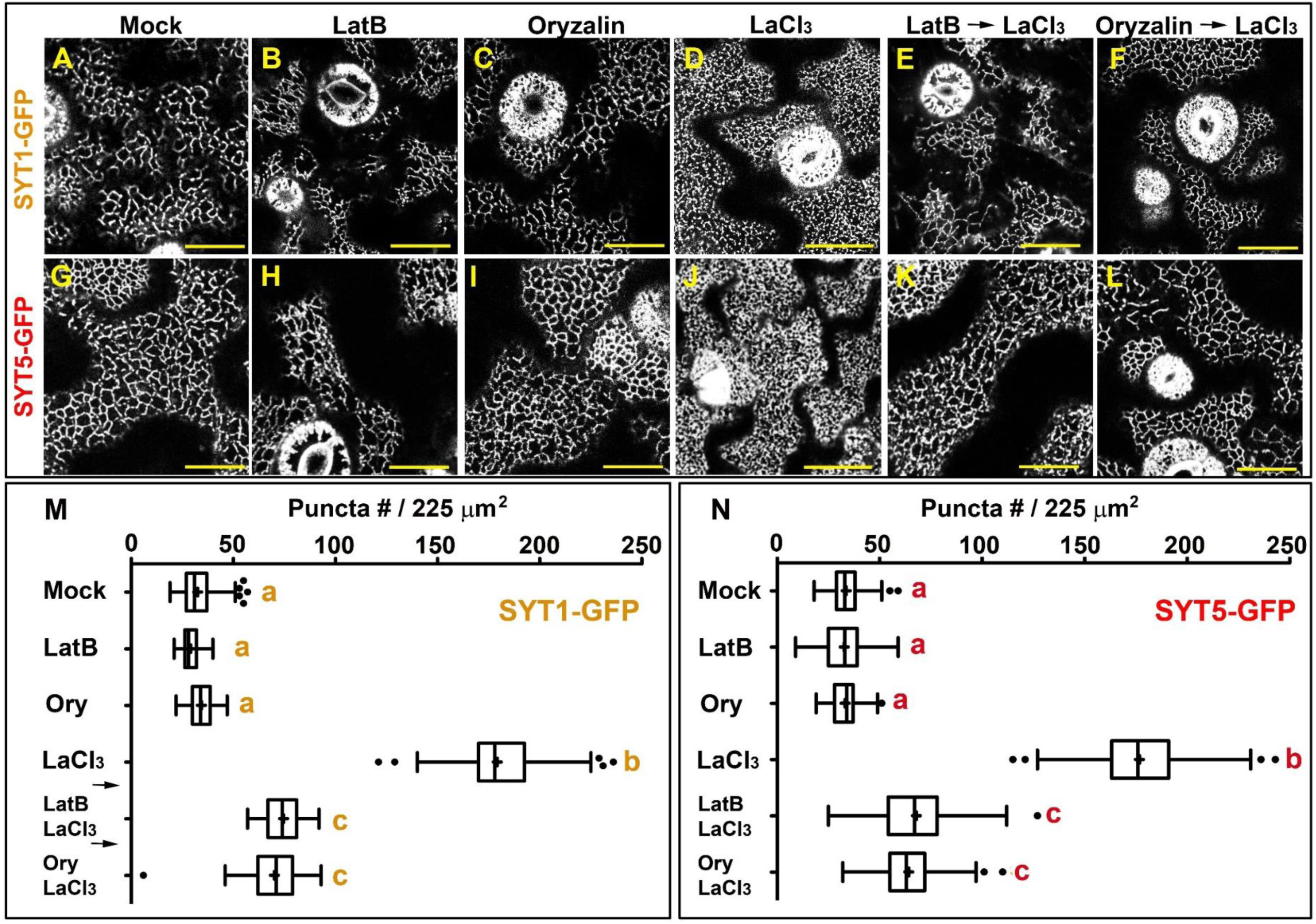
REE-endocytosis induces cytoskeleton-dependent changes in EPCS configuration. Confocal images of the cell cortex in cotyledon epidermal cells expressing the SYT1-GFP (A-F) and SYT5-GFP (G-L) markers. Five-day-old transgenic seedlings grown in one-tenth-MS were transferred to liquid one-tenth-MS, 16 h **(A and G)**, or the same media supplemented with LatB (1 μM, 2 h) **(B and H**), oryzalin (25 μM, 16 h)**(C and I)**, LaCl3 (500 mM, 16 h) **(D and J)** or sequentially treated with LatB (1 μM, 2 h) followed by LaCl3 (500 mM, 16 h) **(E and K**) or oryzalin (25 μM, 16 h) followed by LaCl3 (500 mM, 16 h) **(F-L)** before imaging. **M-N)** Quantification of SYT1-GFP and SYT5-GFP puncta at the cell cortex upon chemical treatments For each treatment, the number of SYT1-GFP and SYT5-GFP puncta was scored from 50-60 arbitrary 225 mm^2^ ROIs from at least 15 cells from 5 different seedlings. In the box and whiskers plots, the center line represents the median number of closed reticules / 225 mm^2^, the top and bottom edges are the 25th and 75th percentiles of the distribution, and the ends of the whiskers are set at 1.5 times the IQR. the minimum / maximum values outside the IQR are shown as outliers (dots). Letters indicate statistically significant differences using Tukey multiple pairwise-comparisons p <0.05 Scale bars = 20 μm

**Figure 7.**
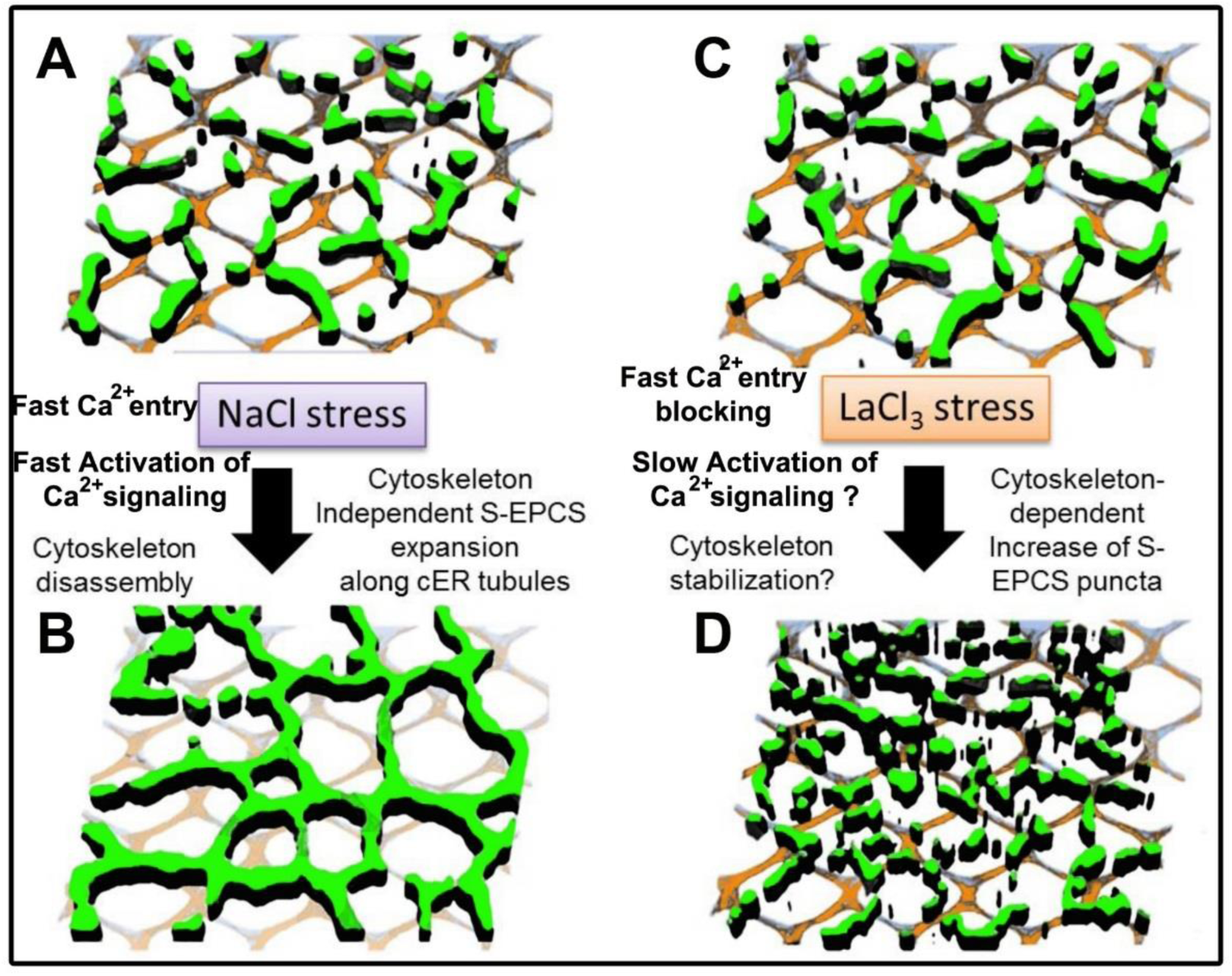
Summary of S-EPCS remodeling mechanisms in response to NaCl and LaCl3 stress. **(A-B)** NaCl stress destabilize the cortical cytoskeleton (orange/blue mesh) and promotes the relocation of SYT1 along the cER tubules effectively promoting cytoskeleton-independent S-EPCS expansions **(C-D**) Long-term treatments with LaCl3 activate endocytosis, increase the activity of the Calmodulin –based sensor GCaMP3, and promote the cytoskeleton-dependent accumulation of SYT1 and SYT5 tethers at the cell cortex effectively increasing the number of S-EPCS.

## Discussion

### Endocytosed REEs as regulators of ER-PM communication

Ca^2+^ is an ubiquitous intracellular secondary messenger involved in many eukaryotic signal transduction pathways **[50-52]**. As high [Ca^2+^]_cyt_ generates toxic intermediates, eukaryotic cells actively compartmentalize Ca^2+^_cyt_ to different organelles and the extracellular space **[53]**. In plants, this process creates up to 10,000-fold [Ca^2+^] differences between the cytoplasm (100 nM) and the apoplast (1 mM), which enables the generation of Ca^2+^ signatures that provide specificity to responses to abiotic, biotic, and developmental stimuli **[54-57]**. Despite the essential role of Ca^2+^ signatures as triggers of signaling cascades, little is known about the elements that regulate the spatio-temporal distribution of Ca^2+^ sensors in plants. Indeed, most of the current information available in plants has been inferred from other eukaryotic systems with very different toolkits of Ca^2+^ channels, transporters, and signaling components **[58]**. In this study, we identify structural and functional similarities between mammalian and plant EPCS tethers, but we also uncover important functional and mechanistic differences of the Ca^2+^ responses mediated by them. Thus, mammalian E-Syts and the plant SSC complex share a common basic organization as both could potentially establish homotypic and heterotypic complexes with their N-terminal domains anchored to the ER, and their C-terminal C2 domains establishing Ca^2+^-dependent interactions with the PM **[28]**. E-Syts and the SSC complex differ, however, in the number of predicted Ca^2+^-responsive C2 domains (two C2s in E-Syt1 **[59]**, and one C2 in the SSC components), and in their noticeably different responses to changes in [Ca^2+^]_cyt_. In mammals, E-Syt1 aggregates and concentrates at membrane junctions following a rise in [Ca^2+^]_cyt_ **[28]**. However, in Arabidopsis, a similar behavior is observed when PM Ca^2+^ channel blockers **[39-42]** /endocytosis inductors **[43, 44]** are used. This differential behavior is difficult to reconcile unless endocytosed REEs were able to trigger intracellular Ca^2+^ signals, effectively offsetting their effect as PM Ca^2+^ channel blockers. Since REEs are internalized via endocytosis upon long-term treatments, and REEs are known to act as allosteric regulators of the activity of Calmodulins **[47]** and C2 containing proteins **[60]**, we propose that endocytosed REEs promote changes in ER-PM communication by either activating Calmodulin signaling or through direct binding to the SSC complex Ca^2+^-binding domains. Still, it is unclear whether EPCS microdomains have an active role in REE endocytosis, following the models proposed in **[61, 62]**, and which molecular components confer specificity to the EPCS tethers’ responses. Given the growing number of EPCS components, Calmodulins, and C2 containing proteins described in Arabidopsis, the elucidation of specific mechanisms underlying REE-induced EPCS reorganization will likely require genetic strategies, such as REEs-resistance screens and/or mutant analyses beyond the scope of this study.

Another functional difference between mammalian E-Syts and plant SYTs is the temporal regulation of their Ca^2+^-mediated responses. In non-excitable mammalian cells, EPCS control intracellular Ca^2+^ levels using store-operated Ca^2+^ entry (SOCE), a fast process that couples the Ca^2+^ influx from the extracellular space to the cytosolic Ca^2+^ release from the ER within seconds **[63]**. These mammalian cells can also sense high [Ca^2+^]_cyt_ and trigger the recruitment, within minutes, of E-Syt1 tethers to SOCE-independent EPCS **[64]**. In contrast, plants lack clear orthologues of the mammalian SOCE components **[58, 65]**, the SYT1-GFP and SYT5-GFP localization changes in response to LaCl_3_ occur within hours, and the Ca^2+^-dependent susceptibility to NaCl and cold stresses is observed days after the stress application **[6, 14]**. Based on these observations, we hypothesize that the plant SSC complex is neither involved in the fast coupling of the extracellular and ER-lumen Ca^2+^ stores nor in the fast responses to [Ca^2+^]_cyt_ changes. Instead, we propose that the observed SSC remodeling in response to REEs are a consequence of the sensing and transduction of sustained Ca^2+^ stress signals that induce long-term cellular adaptive responses, such as the changes in the PM lipid composition previously described for NaCl in **[8]**, and those observed upon REE treatments in this study. Whether the loss of CLB1-GFP signal in response to REEs is a biological consequence of the rearrangement of the SSC complex, or an artifact due to overexpression of CLB1-GFP in a WT background is currently under evaluation.

### Stress-specific regulatory mechanisms controlling S-EPCS organization in Arabidopsis

The cortical ER is a complex arrangement of tubules and small cisternae distributed towards the PM **[66-68]**. EPCS are important substructures within the cER that can be defined as 200-300 nm long and 30 nm in wide cER nanodomains, which anchor to the PM using specialized tethering complexes **[69]**. In a differentiated plant cell, EPCS can be localized in immobile ER tubules **[70]** and areassociated with cortical microtubules **[7, 15, 16, 71]** and the filamentous actin-myosin system **[8]**. Currently, two complementary functions for the cortical cytoskeleton array in EPCS establishment have been proposed. On the one hand, the actin and microtubule networks physically interact with VAP27/NET3C tethering complexes fixing them on specific positions within the cell cortex **[15]**, and this interaction might be required for cargo exchange during endocytic and exocytic trafficking **[16, 71]**. On the other hand, the docking of SYT1-labeled and VAP27-labeled EPCS tethers occurs in microtubule-depleted regions, due in part to the spatial incompatibility between the diameter of microtubules (≈25nm) and the estimated cytosolic gap between the ER and PM at plant, yeast, and mammalian EPCS (≈30 nm) **[69, 72]**. In this report, we identify new commonalities between SYTs and VAP27/NET3C complexes by showing that the SSC complex requires a functional cortical cytoskeleton for proper reorganization in response to REEs. Under these conditions, the SSC and VAP27 complexes could contribute to the overall cER structure and stability by promoting cytoskeleton-dependent EPCS rearrangements in a process likely mediated by Ca^2+^ signals. In this model, the SSC and VAP27 complexes would be enriched, and in close proximity, to regions where the cortical microtubules intersect with the cortical ER–actin network, and the Ca^2+^ and cytoskeleton dependency of the EPCS remodeling would enable the integration of cytoskeleton dynamics with the cER to PM communication required to withstand stress. In response to environmental conditions that induce cytoskeleton disassembly and reorganization (e.g NaCl), an alternative phosphoinositide-associated mechanism would promote cytoskeleton-independent EPCS remodeling and regulate the extent of the cER-PM interaction **[8] (Figure 7**). Together, these examples illustrate the plasticity that governs cER-PM communication, an evolutionary milestone of eukaryotic organisms in their adaptation to environmental stresses.

## Materials and Methods

### Plant materials and growth conditions

*Arabidopsis thaliana* Columbia (Col-0) was used as the wild type and the background for transgenes. Seeds of the mutant alleles *syt5-1* (SALK_036961) and *clb1-2* (SALK_006298) were obtained from the Arabidopsis Biological Resource Center (Ohio State University). Previously published lines in this study are SYT1-GFP **[8]**, GFP-HDEL **[73]**. Plants were grown on half-strength Murashige and Skoog (MS) media (Caisson Labs) or soil (Sunshine mix #4, Sun Gro Horticulture Canada Ltd.) at 22°C with a 16-h light/8-h dark cycle. For the NaCl assays *Arabidopsis* seedlings were grown vertically for 4 d on one tenth-strength MS medium and similar size seedlings were transferred to the same media supplemented with different NaCl concentrations. The root elongation and root hair phenotypes were scored after 9 d.

### Co-immunoprecipitation and large-scale IP for LC-MS/MS

Large-scale immunoprecipitation assays for LC-MS/MS were performed as described before **[74]**, using 5-8 g of 10 day-old Arabidopsis seedlings expressing *p35S::GFP* **[75]** or *pSYT1::SYT1-GFP* **[8]**. For targeted co-IPs, 1 g of 10 day-old Arabidopsis seedlings was frozen in liquid nitrogen. Arabidopsis stable transgenic lines expressing *p35S::GFP* (control),pUB10::CLB1-GFP, and *pSYT5::SYT5-GFP* were used. Total proteins were extracted and immunoprecipitation was performed with GFP-trap beads (Chromotek, Planegg-Martinsried, Germany). Proteins were stripped from the beads by boiling in 50 μl SDS loading buffer for 10 mins. Immunoprecipitated proteins were separated on SDS-PAGE acrylamide gels and western blots were performed using anti-GFP (Santa Cruz Biotechnology sc-9996), anti-SYT1 **[7]**, anti-Mouse IgG-Peroxidase (Sigma A9044), and anti-Rabbit IgG-Peroxidase (Sigma A0545).

### Molecular cloning and generation of transgenic plants

The *CLB1* and *SYT5* constructs used in this study were generated via PCR amplification using the RT-PCR product or genomic DNA as a template and gene specific primers, followed by cloning PCR products into *pENTR/TOPO* (Invitrogen, Carlsbad, CA) or pDONR221. To generate the *pUB10::CLB1-GFP* construct the CLB1 fragment was subcloned into the pB7m24GW,3 vector that contains a 615bp UBQ10 promoter. To generate the *proSYT5::SYT5- GFP* construct, *pEN-L4-proSYT5-R1* and *pEN-L1-SYT5 genomic-L2 and pEN-R2-C-GFP-L3* **[76]** were recombined into *pK7m34GW,0* **[77]** vector. To generate the pCLB1::CLB1-GFP construct the CLB1 pENTR clon was recombined with the destination binary vector pGWB4. All resulting expression vectors were transformed in Arabidopsis via floral dip **[78]**. The selection of transgenic lines was made on half-strength MS medium containing 25 μg/ml hygromycin (pGWB4) or 15 μg/ml glufosinate-ammonium (Sigma-Aldrich) (pB7m24GW,3).

### Chemical Applications

Chemicals were exogenously applied by incubating 5-d-old seedlings in liquid one-tenth-strength of MS medium and supplementing them with 500 µM LaCl_3_ (Sigma-Aldrich), 500 µM GdCl_3_ (Sigma-Aldrich), 5 mM EGTA (Sigma-Aldrich), or 25 μM Oryzalin (Sigma-Aldrich) for 16 h, or with 250µM bis-(*o*-aminophenoxy) ethane-*N, N, N’, N’*-tetra-acetic acid (BAPTA) (Sigma-Aldrich), or 1 µM Latrunculin B (Abcam) for 2 h. The length of the treatments was based on the general toxicity caused by the different chemical compounds in plants.

### Microscopy

#### Image acquisition and quantitative analyses

Living cell images were obtained using a Nikon C1 confocal laser scanning microscope, a Perkin-Elmer spinning disk confocal microscope, and an Olympus FV1000 multiphoton confocal laser scanning microscope. The Nikon C1 confocal laser scanning microscope was equipped with 488 and 515/30nm emission filter and Nikon Plan Apochromat oil immersion objectives (40 × 1.0 NA and 60 × 1.4 NA, respectively). The Perkin Elmer spinning disk confocal microscope was equipped with 488 and 561 nm lasers. The Olympus FV1000 was equipped with 405, 473 and 559 nm lasers and a 60x oil Planon (60 × 1.4 NA). Images were captured using Nikon-EZ C1, Olympus FV1000 and Volocity software, respectively. To quantify the number of “beads” configuration five-day-old Arabidopsis seedlings harboring the SYT1-GFP or SYT5-GFP marker were incubated for 16 h in liquid 1/10-strength MS medium (Mock) or liquid 1/10-strength MS medium supplemented with the different chemicals. For each treatment the number of “beads” labelled by SYT1-GFP or SYT5-GFP in the cortex of cotyledon epidermal cells was scored in at least 50 (15 μm × 15 μm) regions of interest (ROIs) using the cell counter tool of Fiji (ImageJ) (National Institutes of Health, http://imagej.nih.gov/ij/) **[79]**. To compare the fluorescent intensity of the ratiometric CITRINE-1×PHFAPP between control and treated-samples, confocal laser scanning images of 5-d-old epidermal cotyledon cells were acquired from at least 10 individual seedlings. For each data point the fluorescence intensity data was scored from at least 100 (15 μm ×15 μm) ROIs using the integrated density measurement tool of Fiji **[79]**. In this analysis stomatal lineage cells were excluded from the quantification. To compare the fluorescent intensity of the ratiometric GCaMP3 sensor, images of 5-d-old seedlings were acquired using a Nikon stereo microscope SMZ18 equipped with 480/40nm excitation filter, Nikon P2-SHR Plan Apo 0.5x objective and Nikon DS-Ri2 camera. The images were captured using NIS-Elements BR software version 4.60. For each data point the fluorescence intensity data was scored from at least 50 seedlings.In the ratiometric analyses the fluorescent data was normalized using the equation: ΔF/F=(F-F_0_)/F_0_, where F_0_ is the mean intensity of background fluorescence. The data was subject to one way ANOVA analyses to identify statistical differences among treatments. All statistical analyses were performed using the GraphPad Prism 5.0b software.

### Accession numbers

The Arabidopsis Genome Initiative locus identifiers for genes used in this article are SYT1 (*At2g20990*), SYT5 (*At1g05500*), and CLB1 (*At3g61050)*.

## Acknowledgements

This research was undertaken thanks to funding from the NSERC Discovery Grant RGPIN-2019-05568, the Canada Research Chairs program (to A.R.) and the Coordenação de Aperfeiçoamento de Pessoal de Nivel Superior – Brasil (CAPES 88881.189854/2018-01, to B.S.).

