## Supplemental Files for "A putative ER-PM contact site complex relocalizes in response to Rare Earth Elements –induced endocytosis"

### Supplemental Data

| | | Replicate 1 | | Replicate 2 | | Replicate 3 | |
|---|---|---|---|---|---|---|---|
| Gene ID | Protein name | GFP IP | SYT1-GFP IP | GFP IP | SYT1-GFP IP | GFP IP | SYT1-GFP IP |
| - | **GFP** | 25 | 11 | 24 | 18 | 29 | 24 |
| **AT2G20990.1** | **SYT1** | - | 31 | - | 56 | - | 68 |
| **AT1G05500.1** | **SYT5** | - | 11 | - | 23 | - | 39 |
| **AT3G61050.1** | **CLB1** | - | 8 | - | 16 | - | 12 |
| | | Exclusive unique peptide counts | | | | | |

**Table S1: SYT5 and CLB1 are reproducible interactors of SYT1.** The table shows exclusive unique peptide counts corresponding to the indicated proteins in three independent experiments upon immunoprecipitation of SYT1-GFP or free GFP followed by LC-MS/MS analysis.

| Table S2 | SYT1 | SYT5 | CLB1 |
|---|---|---|---|
| Tissue | Total counts number | | |
| Axis of the inflorescence | 4303 | 1196 | 2019 |
| Opened anthers | 1976 | 853 | 1409 |
| Anthers of the mature flower (before opening). | 1214 | 910 | 2065 |
| Anthers of the young flower | 4925 | 2294 | 2487 |
| Carpels of the mature flower (before pollination | 3313 | 1237 | 2209 |
| Carpels of the young flower | 3194 | 981 | 2160 |
| Stamen filaments of the mature flower | 6873 | 2588 | 2976 |
| Petals of the mature flower | 4678 | 1952 | 2330 |
| Sepals of the mature flower | 4519 | 1262 | 1937 |
| Sepals of the young flower | 4261 | 1038 | 2026 |
| Flower 1 | 3082 | 1218 | 1794 |
| Flower 6-8 | 3492 | 1076 | 1790 |
| Flower 9-11 | 4088 | 1234 | 2309 |
| Internode | 5050 | 1528 | 1983 |
| Leaf blade of the mature leaf | 4587 | 821 | 1756 |
| Leaf blade of the young leaf | 5456 | 1333 | 2817 |
| Whole mature leaf | 5029 | 998 | 2081 |
| Petiole of the mature leaf | 4089 | 985 | 1721 |
| Petiole of the senescent leaf | 3112 | 1072 | 1610 |
| Petiole of the young leaf | 6381 | 1716 | 3204 |
| Leaf vein, intermediate 2 | 6839 | 1274 | 2464 |
| Vein of the mature leaf | 3908 | 842 | 1726 |
| Vein of the senescent leaf | 3469 | 987 | 1497 |
| SAM at 7 days after germination | 4058 | 2386 | 2979 |
| Meristem at 10 days after germination | 6011 | 1898 | 3316 |
| Meristem at 11 days after germination | 5992 | 2195 | 3554 |
| Meristem at 12 days after germination | 3769 | 1241 | 2366 |
| Inflorescence meristem at 13 days after germination | 3386 | 1208 | 2086 |
| Inflorescence meristem at 14 days after germination | 4010 | 1092 | 2213 |
| Inflorescence meristem at 15 days after germination | 3839 | 1125 | 2169 |
| Pedicel | 5876 | 1349 | 2578 |
| Pod of the silique 1 | 4872 | 1377 | 2018 |
| Root without apex | 3152 | 1137 | 1391 |
| Root apex | 4968 | 1723 | 1998 |
| Seedling cotyledons | 6650 | 1096 | 2633 |
| Seedling hypocotyl | 6494 | 1246 | 1999 |
| Seedling meristem | 4381 | 2171 | 2189 |
| Seedling root | 4454 | 1580 | 1745 |
| Dry seeds | 8251 | 2270 | 3650 |
| Germinating seeds 1 (first day after soaking) | 4018 | 1318 | 2341 |
| Germinating seeds 2 (second day after soaking) | 6005 | 1509 | 2497 |
| Germinating seeds 3 (third day after soaking) | 6683 | 1502 | 2389 |
| Young seeds 1 | 3370 | 1583 | 2004 |
| Silique 2 | 4066 | 1642 | 2210 |
| Stigmatic tissue | 4085 | 2090 | 1709 |
| **Average read counts** | **4583** | **1425** | **2231** |

**Table S2.** Expression profiles of the SYT1, SYT5, and CLB1 genes across different organs and developmental stages using high-throughput transcriptome sequencing. The values represent the raw number of counts for each gene at a given organ/ developmental stage. Green colors denote higher expression level, yellow colors denote lower expression levels.

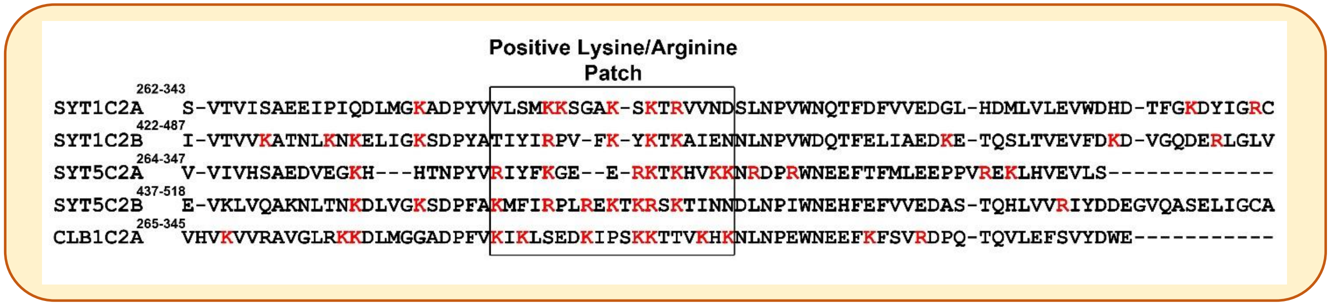


**Figure S1. Multiple sequence alignment of the SYT1/SYT5/CLB1 C2 domains.** The identification of the C2 regions was determine using Pfam and the multiple sequence alignment was generated using Clustal Omega (https://www.ebi.ac.uk/Tools/msa/clustalo/) using seeded guide trees and profile hidden Markov models. The positively charged lysine (K) and arginine (R) aminoacid residues are highlighted in red. The approximate position of the positively charged aminoacid patch is indicated with a rectangle.

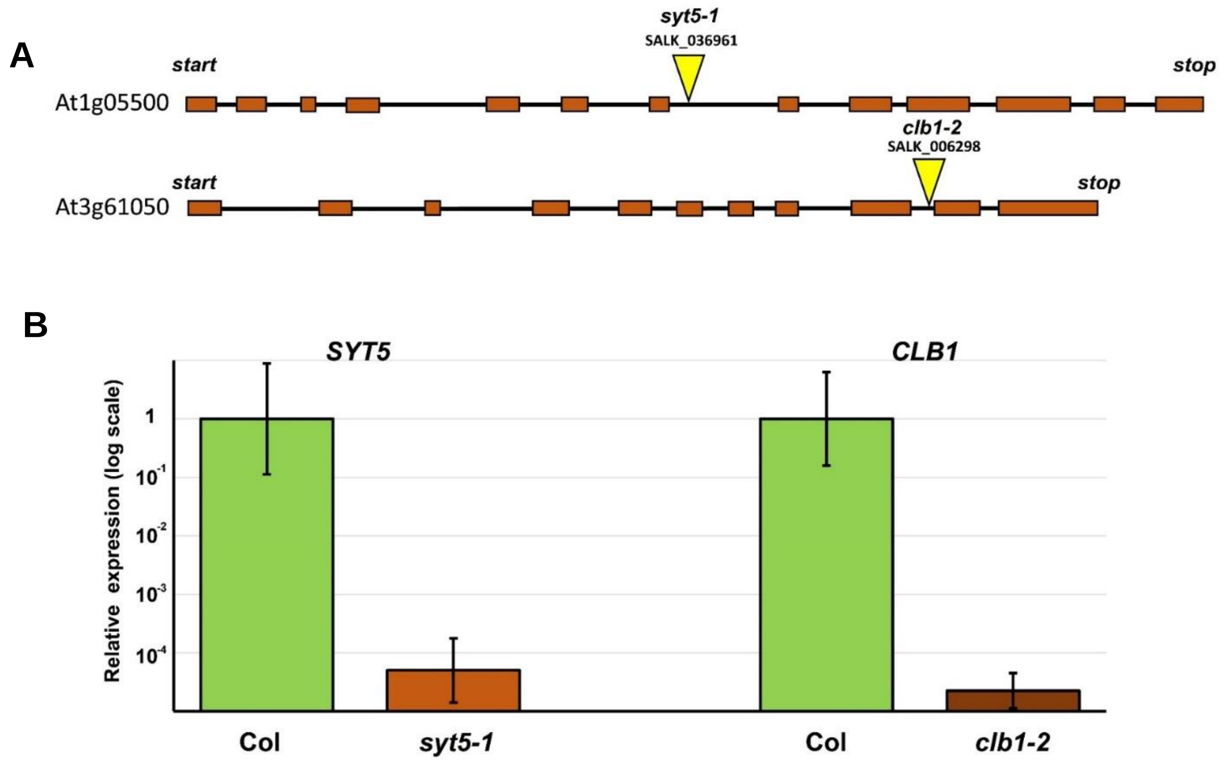


**Figure S2. A)** Exon (orange boxes) – Intron (Black connectors) structured of the At1g05500 and At3g61050 genes. The yellow triangle marks the localization of the T-DNA insertion in the genes. The graph shows the transcript abundance of SYT5 in WT (Col) and *syt5-1* mutant alleles and CLB1 in WT and *clb1-2* mutant alleles measured by qRT-PCR. The levels of expression for SYT5 and CLB1 in mutant backgrounds are reduced more than 10000 fold compared to their controls.

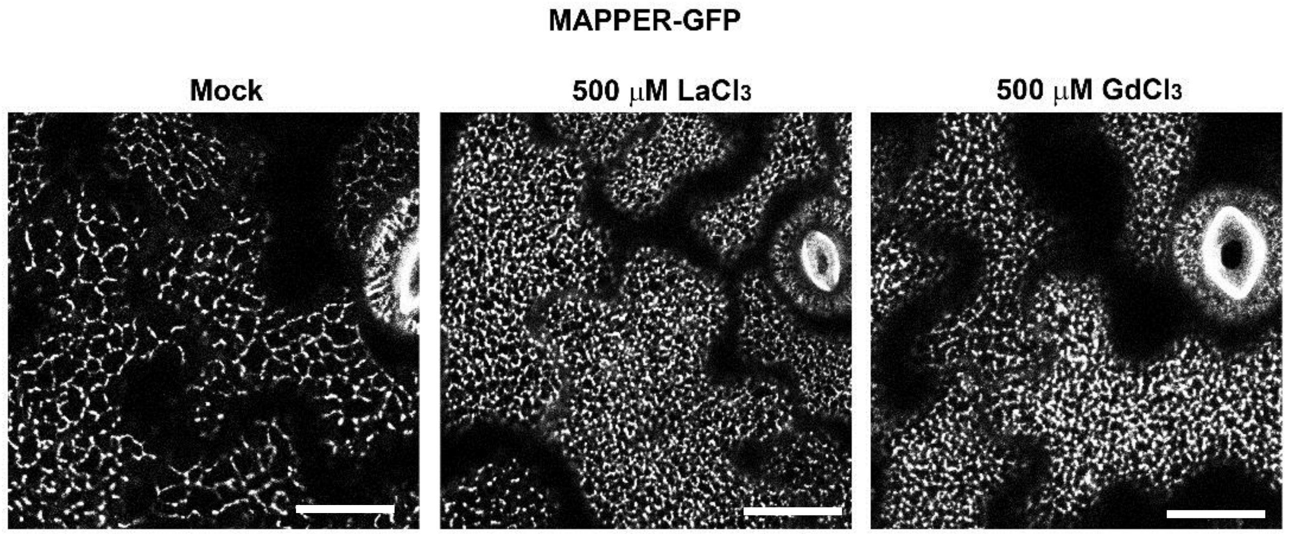


**Figure S3. Treatments with non-selective $Ca^{2+}$ channel blockers increase the number of EPCS at the cell cortex.** 5-d-old MAPPER-GFP seedlings were treated in liquid one-tenth-strength MS medium supplemented with Mock **(A)** $LaCl_3$ (500 µM / 16 h) **(B)** or $GdCl_3$ (500 µM / 16 h) **(C)** before imaging. EPCS accumulation is indicated by the increased number of MAPPER-GFP puncta at the cell cortex upon REE treatment. Scale bars = 20 µm.

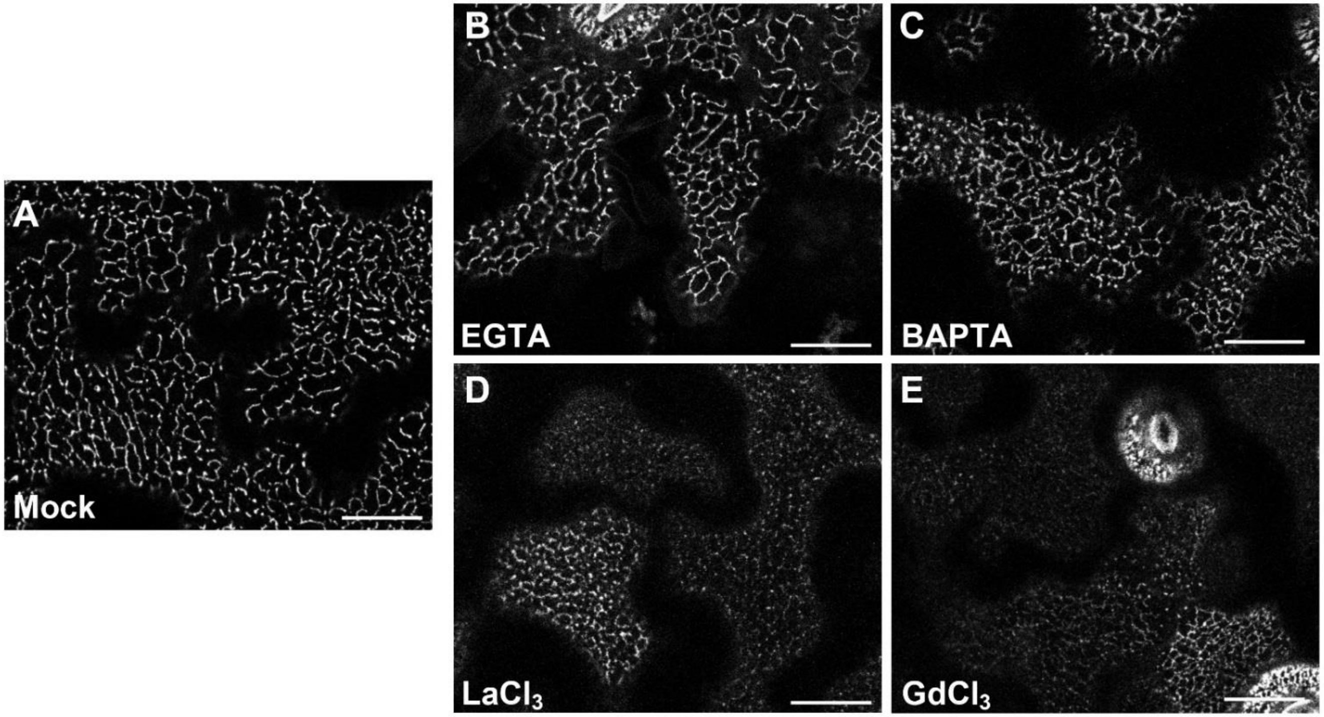


**Figure S4. Treatments with non-selective $Ca^{2+}$ channel blockers reduce the CLB1-GFP signal at the cell cortex.** 5-d-old CLB1-GFP seedlings were treated in liquid one-tenth-strength MS medium supplemented with Mock **(A),** EGTA (5 mM, 16 h) **(B)**, BAPTA (250 mM, 2h) **(C)**, $LaCl_3$ (500 µM / 16 h) **(D)** or $GdCl_3$ (500 µM / 16 h) **(E)** before imaging. The panel shows representative confocal images of the CLB1-GFP signal in cotyledon epidermal cells using similar microscopy settings.

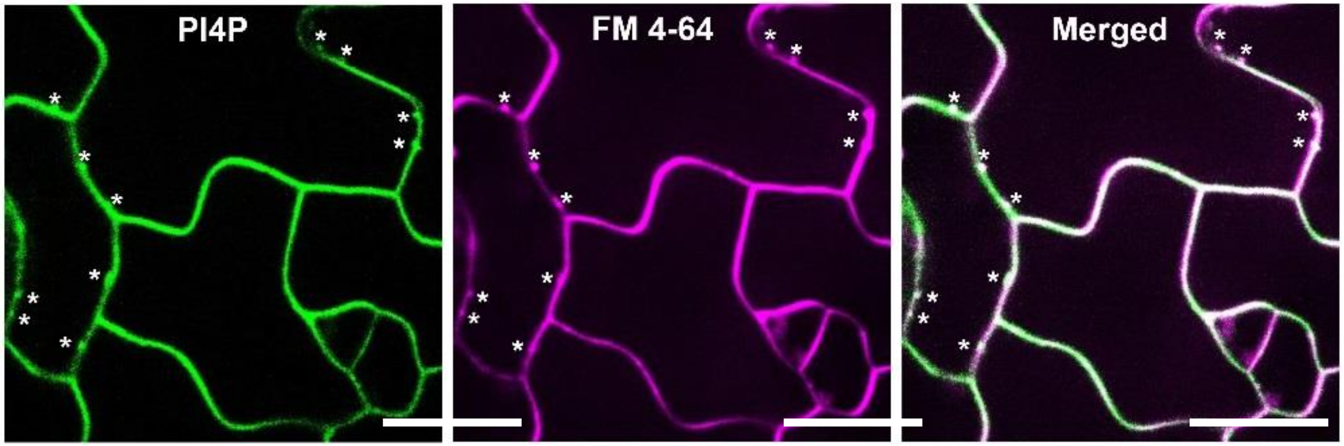


**Figure S5. The CITRINE-1xPH$^{FAPP}$ marker and FM4-64 exhibit overlapping spatial localization patterns at the PM and endocytic vesicles.** A comparison of localization patterns of CITRINE-1xPH$^{FAPP}$ marker and FM4-64 is shown. The localization patterns were examined 180 s after the addition of the FM4-64 dye in 5-d-old cotyledon epidermal cells. Asterisks indicate the position of budding endoytic vesicles. Scale Bars = 20 μm.

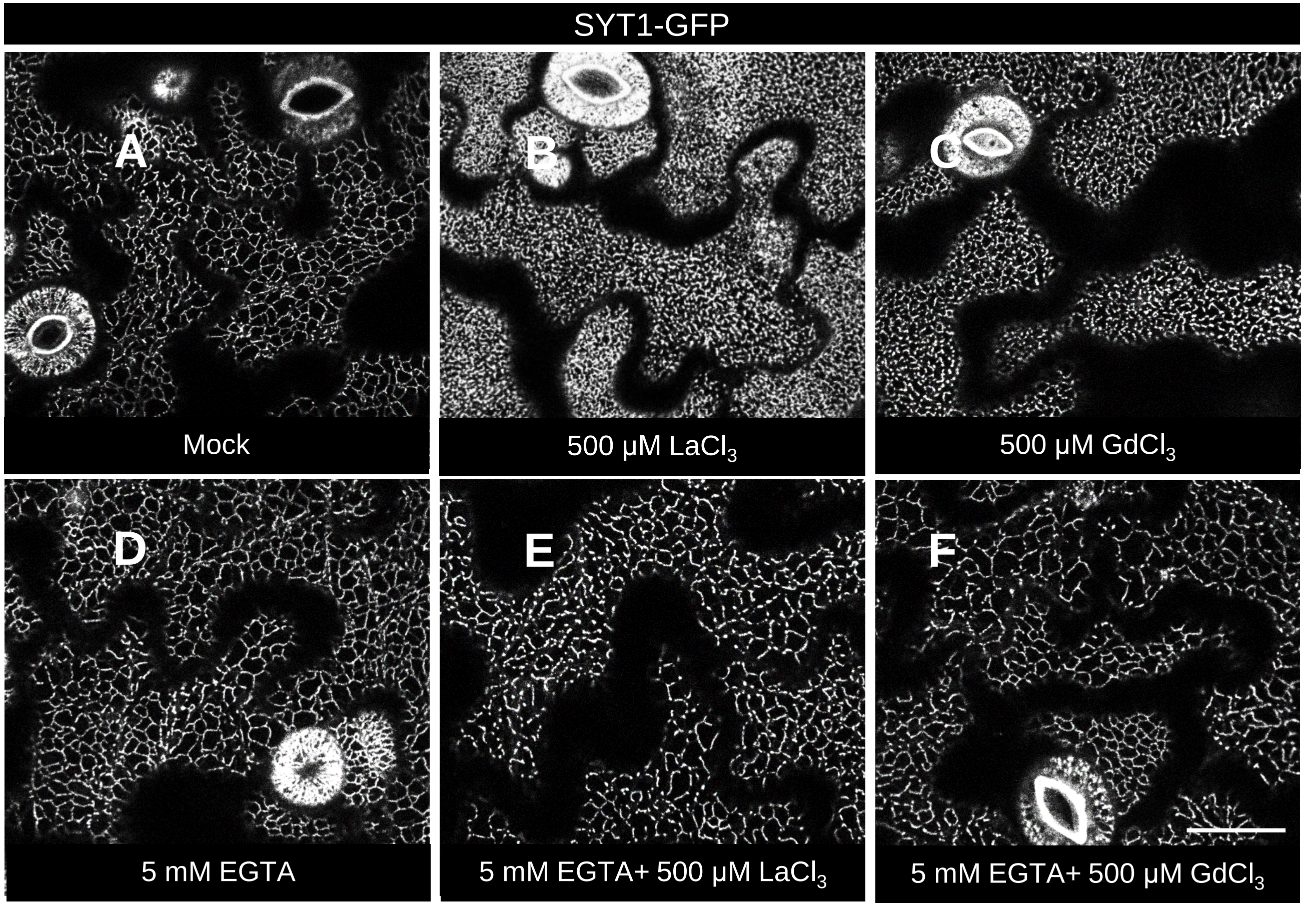


**Figure S6. The addition of EGTA to the growth media inhibits the localization changes associated to REE internalization.** 5-d-old SYT1-GFP seedlings were treated in liquid one-tenth-strength MS medium supplemented with Mock (A) $LaCl_3$ (500 µM / 16 h) (B) $GdCl_3$ (500 µM / 16 h) (C) or the same media supplemented with 5mM EGTA (D-F) before imaging.

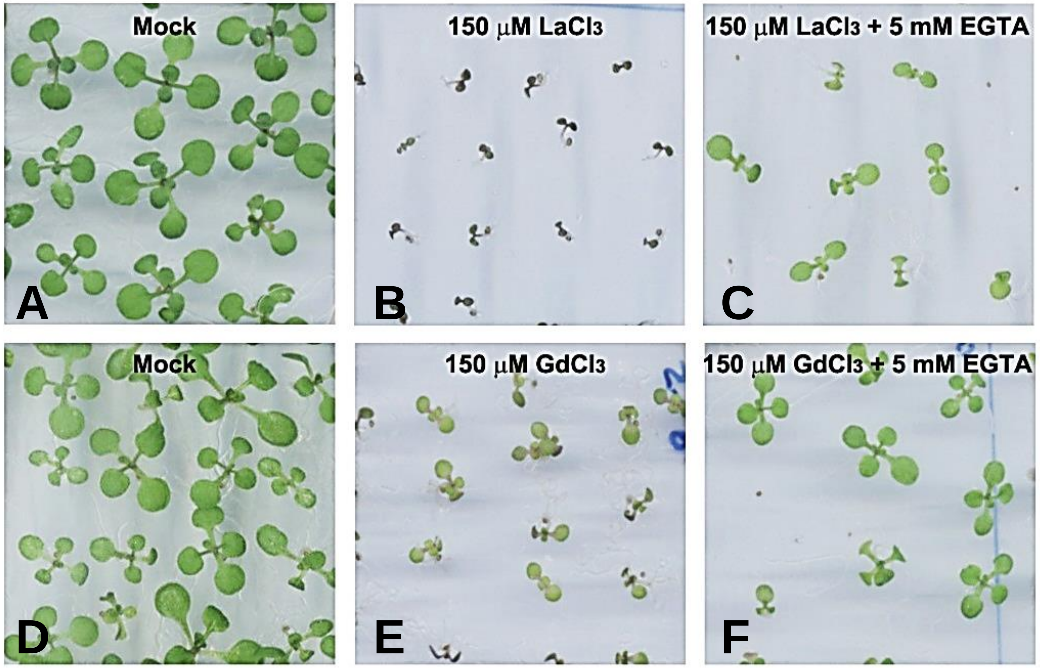


**Figure S7. The exogenous addition of EGTA to the growth media partially reverses the growth in long-term (14 d) REE-treated seedlings.** The images show the development of 14-d-old seedlings grown on one-tenth MS media (A and D) or the same media supplemented with 150 mM $LaCl_3$ (B), 150 mM $LaCl_3$ + 5 mM EGTA (C), 150 mM $GdCl_3$ (E) , or 150 mM $GdCl_3$ + 5 mM EGTA.

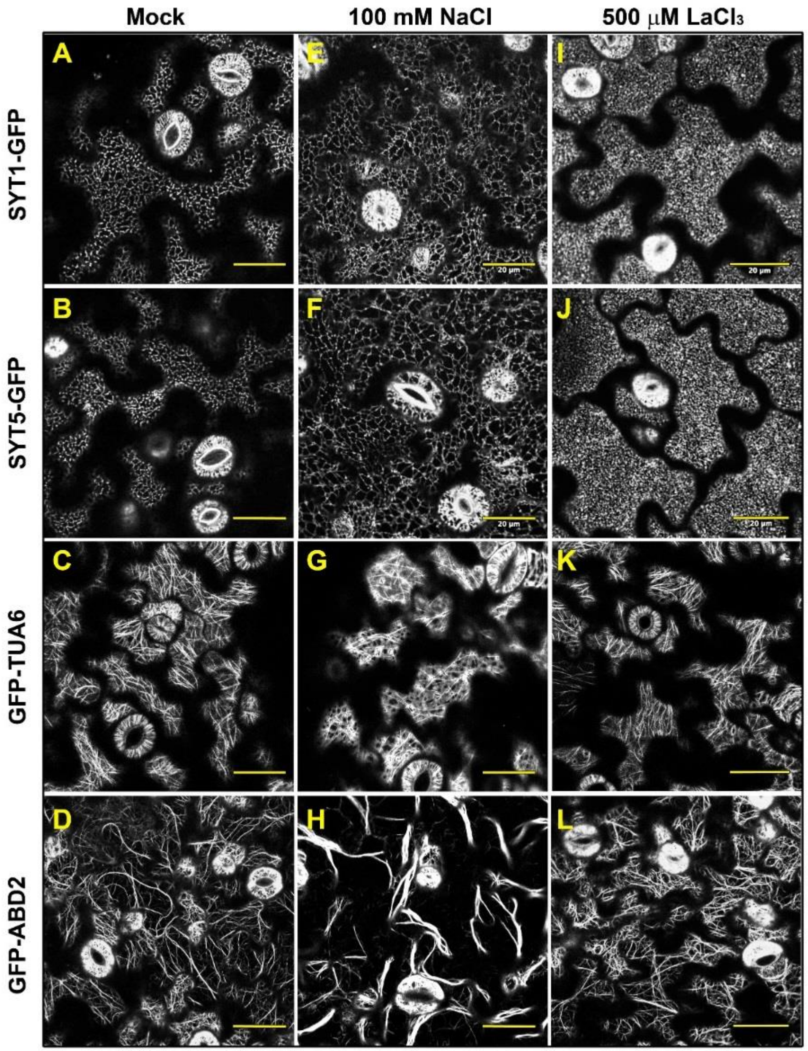


**Figure S8. Effect of NaCl and $LaCl_3$ treatments on SYT1-GFP and SYT5-GFP relocalization and cortical cytoskeleton organization.** Representative images of the SYT1-GFP and SYT5-GFP S-EPCS markers, GFP-TUA6 microtubule marker, and GFP-ABD2 actin filaments marker upon treatments with mock (A-D), 100 mM NaCl, 16 h (E-H) or 500 µM $LaCl_3$, 16 h (I-L). The images were taken in 5-d-old cotyledon epidermal cells. The effect of the 100 mM NaCl stress was characterized by SYT1-GFP and SYT5-GFP signal expansion along cortical ER tubules, microtubule depolymerization, and actin filaments bundling. The effect of $LaCl_3$ was characterized by an increase in the number of cortical SYT1-GFP and SYT5-GFP structures without gross effects in the cortical microtubules or actin filaments organization. Scale bars (A-L) = 20 μm.

#### Table S3. Primers used in this study

| Primer ID | Sequence (5’ to 3’) |
| --- | --- |
| CLB1_CDS F | CACCATGGGTTTGATTTCTGGGATTCTG |
| CLB1_CDS R | CTGCTGTTTTGCACCATCGTTTTCTGG |
| CLB1_ PROM | CACCCTAATATTATGCACGCTT |
| AttB1_MAPPER | GGGGACAAGTTTGTACAAAAAAGCAGGCTATGGATGTATGCGTCCGTCTTGCCCTGT |
| AttB2_MAPPER | GGGGACCACTTTGTACAAGAAAGCTGGGTTCAAGTTACTGAATCTTTCTTCTTCC |
| SYT5_CDS_F | GGGGACAAGTTTGTACAAAAAAGCAGGCTCCATGGGTTTCATAGTCGGCGTTGTAATCG |
| SYT5_CDS_R | GGGGACCACTTTGTACAAGAAAGCTGGGTCGGAATCACGATAAATTGATTGAGC |
| SYT5_PROM | CACCAAGAAAGCGTGATGGCAAAGCCA |
